# An unbiased high-throughput drug screen reveals a potential therapeutic vulnerability in the most lethal molecular subtype of pancreatic cancer

**DOI:** 10.1101/791913

**Authors:** Chun-Hao Pan, Yuka Otsuka, BanuPriya Sridharan, Melissa Woo, Sruthi Babu, Cindy V. Leiton, Ji Dong K. Bai, David K. Chang, Andrew Biankin, Louis Scampavia, Timothy Spicer, Luisa F. Escobar-Hoyos, Kenneth R. Shroyer

**Affiliations:** Department of Pathology, Renaissance School of Medicine, Stony Brook University, Stony Brook, NY, USA; Molecular and Cellular Biology Graduate Program, Brook University, Stony Brook, NY, USA; The Scripps Research Institute, Florida Campus, Jupiter, FL, USA; In vitro In vivo Translation, GSK, Collegeville, PA, USA; Simons Summer Research Program, Stony Brook University, Stony Brook, NY, USA; Department of Family, Population & Preventive Medicine, Renaissance School of Medicine, Stony Brook University, Stony Brook, NY, USA; Wolfson Wohl Cancer Research Centre, Institute of Cancer Sciences, University of Glasgow, Glasgow, UK; West of Scotland Pancreatic Unit, Glasgow Royal Infirmary, Glasgow, UK; David M. Rubenstein Center for Pancreatic Cancer Research, Memorial Sloan Kettering Cancer Center, New York, US; Genetic Toxicology and Cytogenetics Research Group, Department of Biology, School of Natural Sciences and Education, Universidad del Cauca, Popayán, Colombia

**Keywords:** Pancreatic ductal adenocarcinoma, Keratin 17, Predictive biomarker, Chemoresistance, Drug screen, Combined therapy

## Abstract

Pancreatic ductal adenocarcinoma (PDAC) is predicted to become the second leading cause of cancer-related deaths in the United States by 2020, due in part to innate resistance to widely used chemotherapeutic agents and limited knowledge about key molecular factors that drive tumor aggression. We previously reported a novel negative prognostic biomarker, keratin 17 (K17), whose overexpression in cancer results in shortened patient survival. In this study, we aimed to determine the predictive value of K17 and explore the therapeutic vulnerability in K17-expressing PDAC, using an unbiased high-throughput drug screen. Patient-derived data analysis showed that K17 expression correlates with resistance to Gemcitabine (Gem). In multiple *in vitro* models of PDAC, spanning human and murine PDAC cells, we determined that the expression of K17 results in a more than two-fold increase in resistance to Gem and 5-fluorouracil, key components of current standard-of-care chemotherapeutic regimens. Furthermore, through an unbiased drug screen, we discovered that Podophyllotoxin (PPT), a microtubule inhibitor, showed at least two-fold higher sensitivity in K17-expressing compared to K17-negative PDAC cells. In the clinic, another microtubule inhibitor, Paclitaxel (PTX), is used in combination with Gem as a first line chemotherapeutic regimen for pancreatic, breast, lung, and ovarian cancer. Surprisingly, we found that when combined with Gem, PPT but not PTX, was synergistic in inhibiting the viability of K17-expressing PDAC cells. This provides evidence that PPT or its derivatives could potentially be combined with Gem to enhance treatment efficacy for the approximately 50% of PDACs that express high levels of K17. In summary, we reported that K17 is a novel target for developing a biomarker-based personalized treatment for PDAC.

## 1. Introduction

Pancreatic Ductal Adenocarcinoma (PDAC) is one of the deadliest types of cancer, with a 5-year survival rate at only 7%^1^. This extremely poor prognosis is due in part to the lack of effective screening strategies to detect PDAC at early stages^2^, and to the inherent resistance of these tumors to currently available first-line chemotherapeutic agents^3,4^.

The majority of patients are diagnosed at late metastatic or advanced stages and as a result, only 10-15% of PDACs are candidates for surgical resection^5^. With or without surgery, patients are subjected to chemotherapy where Gemcitabine (Gem) was the baseline treatment for more than a decade and is still employed^6^. Currently, Gem, combined with paclitaxel albumin-bound nanoparticles (nab-PTX, Abraxane)^7^ and the FOLFIRINOX regimen (5-Fluorouracil [5-FU], leucovorin, irinotecan, and oxaliplatin)^8^ are two standard-of-care therapies that have been found to improve the survival rates compared to the use of Gem alone^9^. However, despite this improvement, PDACs usually show only weak response to all current treatment regimens^10^. Therefore, exploring biomarker-driven novel therapies is a critical priority for improving therapeutic outcomes for PDAC patients.

To this end, it was reported and recently validated in several cohorts that PDAC is composed of two molecular subtypes, based on distinct gene expression profiles that impact on patient survival^11–14^. These differential gene expression signatures are now being used as biomarkers for molecular subtyping^15–17^, however, whether the proteins encoded by these genes cause resistance to treatment and if they could be exploited as therapeutic targets for PDAC remain unexplored.

We independently demonstrated that keratin 17 (K17), one signature gene overexpressed in the basal-like PDAC^12^, is an independent negative prognostic biomarker and is as accurate as molecular subtyping to predict PDAC patient survival^15^. However, it is still unknown if K17 is predictive for treatment response and if K17 expression in PDAC cells sensitizes them to specific and currently available small-molecule inhibitors. To address these questions, we evaluated K17 mRNA expression of PDACs from patients that received Gem treatment as standard-of-care versus no treatment. This led us to identify that K17 is a predictive marker of Gem response. In addition, a high-throughput small-compound screen uncovered a novel therapeutic vulnerability of tumors bearing K17 expression. Our findings support the conclusion that K17 could be a target for the development of a novel, and potentially more effective biomarker-based personalized therapy for PDAC.

## 2. Methods

### 2.1. Predictive-value analyses from patient-derived samples

K17 mRNA expression levels of PDAC cases were acquired from the Australian Pancreatic Cancer Genome Initiative (APGI)^13^. K17 mRNA expression and survival were evaluated in 94 PDAC patients that were treated with adjuvant Gem alone or received no treatment. Based on the established cutoff of the maximum likelihood fit of a Cox proportional hazard model^15^, we applied the 76^th^ percentile of mRNA expression to categorize patients into high-K17 versus low-K17 groups. Overall survival in high-versus low-K17 mRNA was determined using the Kaplan-Meier method, calculated from the date of diagnosis to the date of death. Patients still alive at the last follow-up were censored. Prognostic and predictive analyses were performed based on the criteria described by Ballman^18^. Adjusting for potential confounders, a multivariate analysis was performed by Cox proportional hazard regression. Statistical significance was set at p<0.05 and analysis was done using SAS 9.4 (SAS Institute) and GraphPad Prism 7 (Graph Pad Software).

### 2.2. Compounds tested

Gem (purity > 99%), 5-FU (purity > 99%), Podophyllotoxin (PPT, purity > 99%), Taxol (PTX, purity > 95%), Mitoxantrone (purity > 99%), and Tyrophostin AG 879 (purity > 99%) were purchased from Sigma-Aldrich (St. Louis, MO, USA). These drugs were dissolved in 100% dimethyl sulfoxide (DMSO, Fisher BioReagents) with a stock concentration of 20 mM and were prepared for cell experiments at final DMSO concentrations at 0.1%.

### 2.3. Cell culture

Human L3.6 PDAC cell line (Kras_G12A_)^19^ was a gift from Dr. Wei-Xing Zong (Rutgers University). MIA PaCa-2 PDAC cells (Kras_G12C_, p53R_248W_) were obtained from American Type Culture Collection. Murine Kras_G12D_, p53_R172H_ pancreatic cancer cells (KPC) cell line was a gift from Dr. Gerardo Mackenzie (University of California at San Diego). Cells were cultured at 37 °C in a humidified incubator under 5% CO_2_ in Dulbecco’s Modified Eagle Medium (DMEM, Gibco) supplemented with 10% fetal bovine serum (FBS, ThermoFisher) and 1% penicillin and streptomycin (P/S, Gibco).

### 2.4. CRISPR-Cas9 mediated K17 knockout in PDAC cell line

CRISPR-Cas9 mediated knockout cell pool of K17 (KRT17-gene name) in L3.6 cells were generated by Synthego Corporation (Redwood City, CA, USA). To generate these cells, ribonucleoproteins containing the Cas9 protein and the synthetic chemically modified single-guide RNA (CCAGTACTACAGGACAATTG) were electroporated into the cells using Synthego’s optimized protocol (https://www.synthego.com/resources/all/protocols). The genetic editing efficiency was assessed upon recovery (48 hours post electroporation). Genomic DNA was extracted from the cells, PCR amplified and sequenced using Sanger sequencing. The resulting chromatograms were processed using Synthego Inference of CRISPR edits software (ice.synthego.com).

### 2.5. Overexpression of K17 in in PDAC cell lines

MIA PaCa-2 and KPC cells were stably transduced to express either empty vector (EV) or human K17 as K17 gain-of-function (GOF) cell line models. Briefly, cells were transduced for 24 hours in medium supplemented with 10% FBS for 24 hours, followed by fluorescence-activated cell sorting.

### 2.6. Protein extraction and western blot analysis

Cells were harvested using RIPA buffer containing protease and phosphatase inhibitor cocktail (ThermoFisher Scientific). The lysates were sonicated, centrifuged at 13,000 rpm for 10 minutes, and the supernatant was collected. The protein concentration of the cell lysates was measured using a Bradford protein assay kit (Bio-Rad) according to the manufacturer’s instructions. Equal amounts of proteins were separated by 12% SDS polyacrylamide gel electrophoresis. Immunoblotting was performed with primary antibodies to K17^20–22^ (a gift from Dr. Pierre Coulombe, University of Michigan) and GAPDH (Cell Signaling Technology), followed by infrared goat anti-mouse or goat anti-rabbit IgG secondary antibodies (LI-COR Inc.). Western blot images were captured by LI-COR Odyssey Imaging machine and images were quantified using Image Studio Lite software (LI-COR Inc.).

### 2.7. Immunofluorescence imaging

Cells were first fixed in ice-cold methanol for 5 minutes at 20 °C, permeabilized with 0.25% Triton X-100 for 10 minutes at room temperature and blocked in 10% donkey serum (Sigma-Aldrich) dissolved in PBS (Gibco) for 1 hour. Primary K17 antibody^20^ diluted in 10% donkey serum was incubated overnight. Fluorescence conjugated goat anti-rabbit secondary antibody (Abcam) was incubated at dark for 1 hour. Cells were mounted with Vectashield (Vector laboratories) with DAPI.

### 2.8. The Library of Pharmacologically Active Compounds (LOPAC) drug screening

The library currently available at Scripps was purchased from Sigma-Aldrich. Two screens were performed in 1536 well format, which include a screen performed in L3.6 K17-expressing cells and counterscreen in L3.6 K17 knockout (KO) cells. Cell viability was examined using the CellTiter-Glo^®^ Luminescent Cell Viability Assay (Promega), a homogeneous method to determine the number of viable cells in culture based on quantitation of the ATP present, which signals the presence of metabolically active cells. First, 500 cells per well in a 5uL volume were plated (Greiner part # 789173) in DMEM (10% FBS + 1% P/S) and incubated overnight at 37C 5% CO2, and then compounds were added at 2 μM final concentration followed by an additional incubation for 48 hours. Next, 5 μL of CellTiter-Glo^®^ reagent was added to all wells. After incubation at room temperature for 10 minutes to ensure stabilization of luminescence, the plate was read by ViewLux™ (PerkinElmer). Raw assay data was analyzed using Symyx software (Symyx Technologies Inc.). Activity of each compound was calculated based on Average + 3 standard deviation using the following equation:

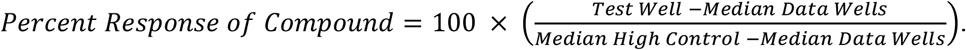

Test Well = Cells + compound, Data Wells = All test wells, High Control = media only plus DMSO. A mathematical algorithm was used to determine active compounds (Hits). Two values were calculated: (1) the average percent response of all compounds, and (2) three times their standard deviation. The sum of these two values was used as a cutoff parameter, i.e. any compound that exhibited greater percent activation than the cutoff parameter was declared active as a hit (p<0.0001).

### 2.9. Cell viability and proliferation assays

Cell viability was examined using the CellTiter-Glo^®^ Luminescent Cell Viability Assay (Promega), or the WST-1 Cell Proliferation Reagent (Sigma-Aldrich). Cells were plated in 96 well plate at 6 × 10^4^ cells/well and incubated overnight, drugs were added with 10-point dose-response titrations in triplicate (0.5 - 20000 nM) or at indicated concentration for 48 hours. For WST-1 colorimetric assay, 10 uL of WST-1 was added per well, and the plate was incubated for 30 minutes and gently shaken. The absorbance was measured using a microplate (ELISA) reader at 450 nm and the reference at 630 nm. To measure cell proliferation, cells were plated at 4 × 10^4^ cells/well in 96-well plates and the relative proliferation index was measured by WST-1 assay at each time point.

### 2.10. Flow cytometric cell cycle analysis

The percent cells at each cell cycle phase was assessed through propidium iodide (PI; Sigma-Aldrich) nuclear staining. Cells were seeded at 3 × 10^5^ cells/dish in 60 mm dishes for 24 hours, and were exposed to DMSO or PPT treatment at 20 nM for 48 hours. After treatments, cells were harvested and stained with PI in Kreshan modified buffer for 30 minutes. The percent cells at each cell cycle phase and apoptosis were measured by flow cytometry (BD Biosciences) and data were analyzed by ModFit LT™ (Verity Software House).

### 2.11. Statistics

The half maximal inhibitory concentration (IC50) of single drug treatment was determined by specificity of compounds was calculated by: 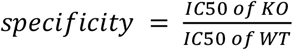, and defined as following: less than 0.5 to 2: medium, greater than 2: high. The combination index (CI) and IC50 for drug combinations were calculated using CompuSyn software (www.combosyn.com), according to the Chou-Talalay model, one of the most widely used methods for detecting and quantifying synergistic interactions between drugs^23^. CI<0.9 indicates a synergistic effect. CI=0.9-1.2 indicates an additive effect. CI>1.2 indicates an antagonistic effect. The statistical significance between two groups was determined using Student’s *t*-test. Data were expressed as means ± standard deviation (SD) or standard error mean (SEM), and *p<0.05, **p<0.01, ***p<0.001 were considered significant.

## 3. Results

### 3.1. K17 is a novel predictive biomarker of Gemcitabine chemotherapy in PDAC

We previously reported that K17 is a prognostic biomarker for PDAC patients. Here, we set out to test if K17 could be a predictive biomarker in PDAC. Using APGI patient data^13^, we first defined K17 status in PDACs by analyzing K17 mRNA expression and applying the previously established cut-off^15^ to categorize high- and low-K17 cases (K17 mRNA Z-scores ranged from −0.66 to 11.17). High-K17 cases were defined as those in the top 24^th^ percentile of K17 expression, while low-K17 cases were the lowest 76^th^ percentile (Fig. 1A). This 76^th^ threshold, which provides maximal stratification of survival differences based on K17 mRNA expression, was trained and subsequently adjusted previously. We first validated the prognostic value of K17. Patients with high-K17 PDACs had a shorter median survival (14 months) than patients with low-K17 PDACs (23 months) (HR = 1.8, log-rank *P* = 0.0334) (Fig. 1B). Furthermore, the prognostic value of K17 mRNA status was independent of pathological stage (Fig 1C).

**Figure 1.**
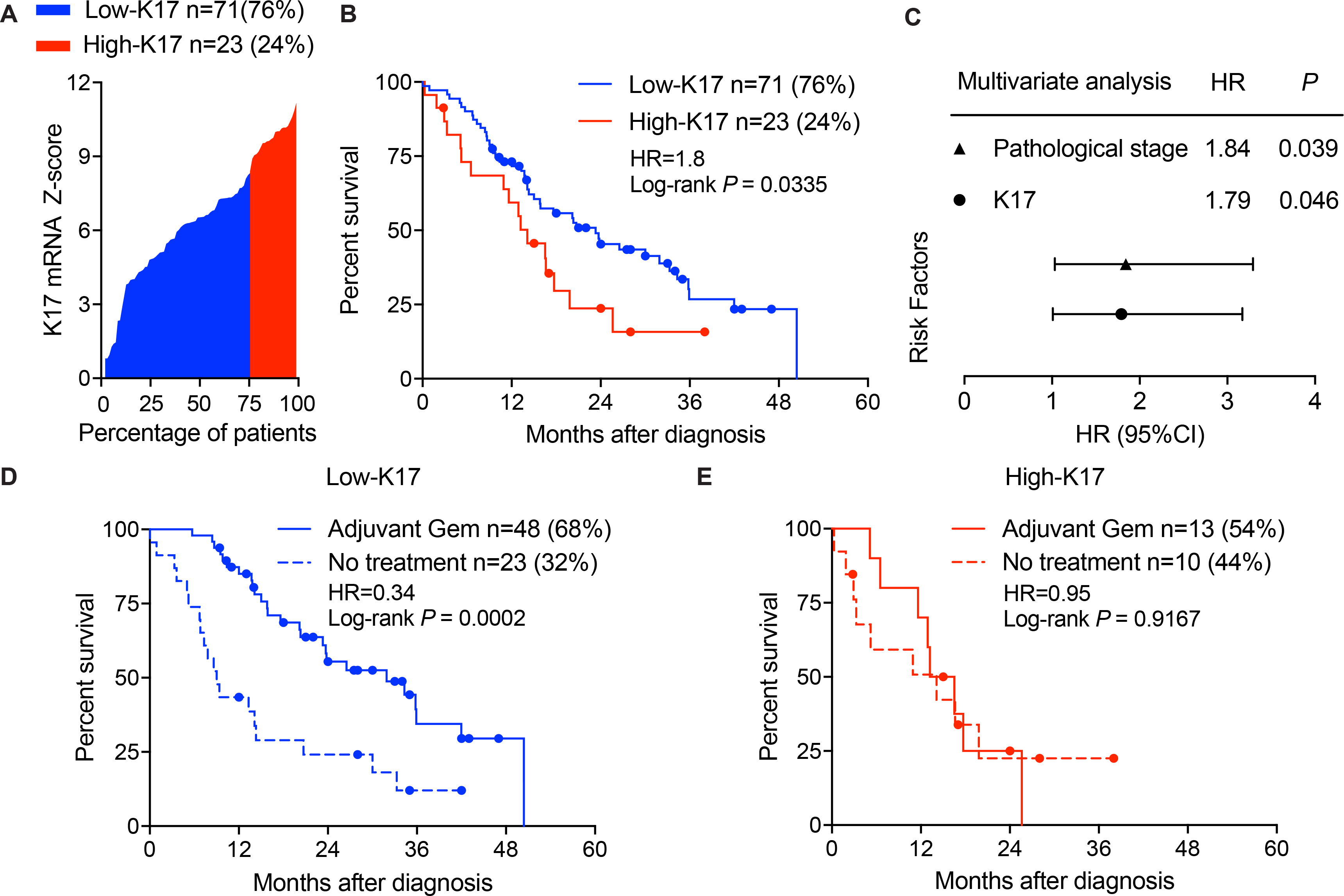
K17 predicts response of Gemcitabine in PDAC patients. **A.** K17 mRNA expression level of patient samples from the APGI cohort are shown in a waterfall plot. The established cut-off of 76^th^ percentile^15^ was applied to categorize patients into high- and low-K17 groups. 76% of PDAC cases below cut-off were classified as low-K17 (blue), and 24% of cases above the cut-off were defined as high-K17 (red). **B.** Kaplan-Meier curves show the overall survival of patients with high-K17 and low-K17 PDACs. Hazard ratios (HR) and log-rank p-value are shown. **C.** Forest plot shows the multivariate analysis from risk factors of the pathological stage and K17 mRNA as a binary variable. Pathological stage and K17 show significant p-values. **D.** Patients with low-K17 PDACs exhibited significantly longer survival after adjuvant Gem therapy, compared with those who did not receive Gem. Hazard ratios (HR) and log-rank p-value are shown. **E.** Patients with high-K17 PDACs exhibited no overall survival differences between groups with or without Gem treatment. Hazard ratios (HR) and log-rank p-value are shown.

To determine if K17 expression predicts response to treatment with Gemcitabine (Gem), we analyzed the survival of patients harboring high or low K17-expressing tumors, comparing treatment of Gem alone (adjuvant Gem) or no treatment using the APGI data. The low-K17 group treated with adjuvant Gem had a median survival of 32 months, which was significantly longer than the no treatment group, with a median survival of 9 months (HR = 0.34, log-rank *P* = 0.0002) (Fig. 1D). In contrast, the high-K17 PDACs did not show differences in the median survival between adjuvant Gem and no treatment groups (HR = 0.95, log-rank *P* = 0.9167) (Fig. 1E). This shows that the low-K17 group responded to Gem treatment but high-K17 group did not show any treatment response, suggesting that there was a qualitative predictive interaction according to Ballman *et al*^18^. Thus, beyond its role as a prognostic biomarker for PDAC, K17 is predictive for the response to Gem.

### 3.2. K17 actively drives chemoresistance to first-line therapeutic agents

Given an association between K17 expression and resistance to Gem, we set out to identify whether K17 expression actively promotes resistance to Gem and another key chemotherapeutic agent 5-Fluorouracil (5-FU), which are major components for the two first-line chemotherapies. To test this, we first generated a K17 loss-of-function (LOF) human cell line model, to assess the response to Gem and 5-FU. Using CRISPR-Cas9 technology, we knocked out endogenous expression of K17 from L3.6 human PDAC cell line (Fig. 2A-B). Isogenic cells with and without expression of K17 were treated with Gem or 5-FU and the IC50 was determined. We found that K17-expressing cells showed significantly higher IC50 values of Gem (Fig. 2C, 2-fold) and 5-FU (Fig. 2D, 2-fold) compared with K17 knockout (KO) cells.

**Figure 2.**
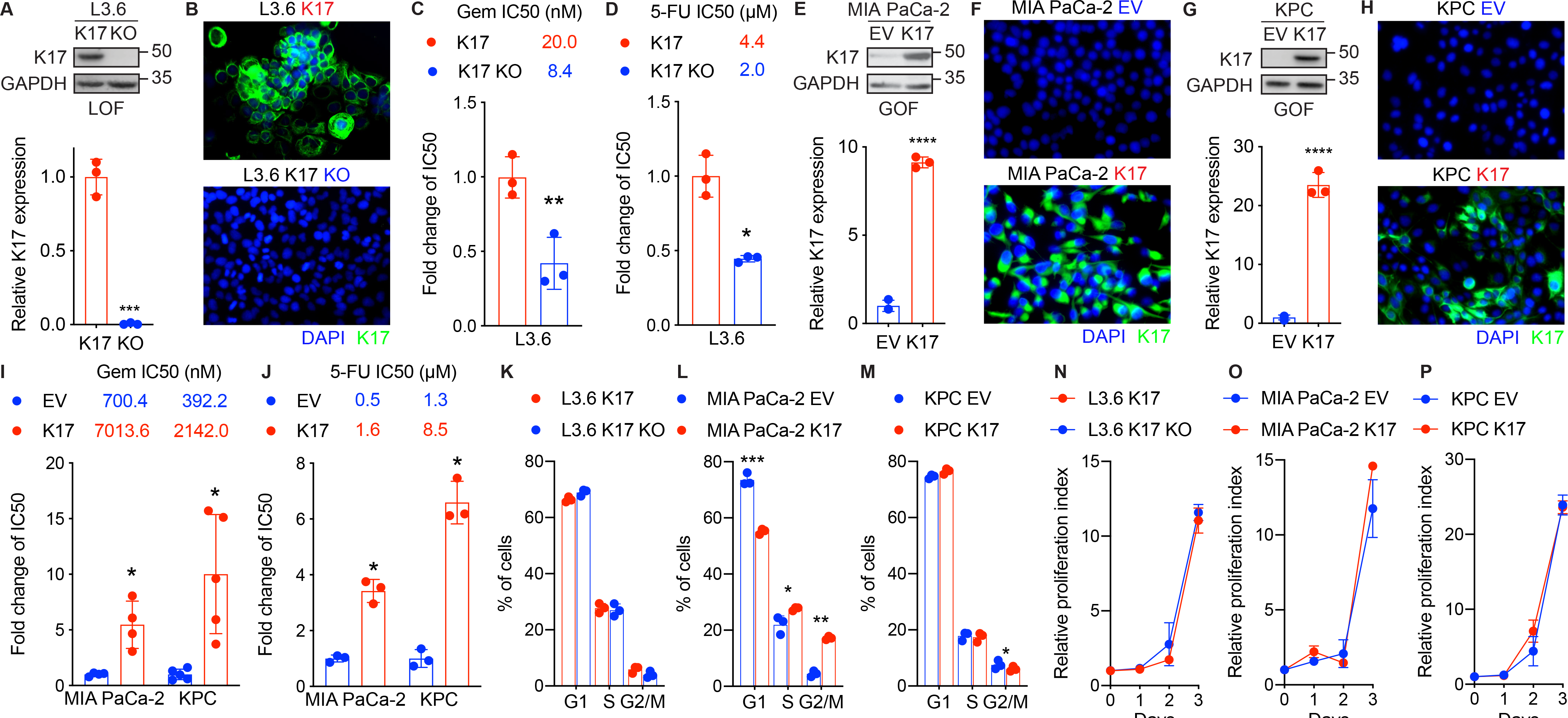
K17 causes chemoresistance to Gem and 5-FU in PDAC. **A.** Genetic manipulation of K17 expression to generate stable cell line model. K17 loss-of-function (LOF) model: L3.6 human PDAC cell line expresses endogenous K17. CRISPR-Cas9 technique was used to generate K17 knock out (KO) cells. Western blot and quantification are shown. **B.** Immunofluorescence images of L3.6 cell line model. Filament form of K17 is shown in green, and the nucleus is indicated by DAPI stain in blue. **C-D.** L3.6 cells expressing K17 showed significantly higher IC50 value of Gem (C) and 5-FU (D) than cells had K17 KO. IC50 values and fold change of IC50 are shown. **E.** Generation of human K17 gain of function (GOF) cell line model. MIA PaCa-2 PDAC cell line express low level of K17. Cells were transduced to stably express either EV (empty vector) or human K17 (K17) as a human K17 GOF model. Western blot and quantification are shown. **F.** Immunofluorescence images of MIA PaCa-2 cell line model. Filament form of K17 is shown in green, and the nucleus is indicated by DAPI stain in blue. **G.** Generation of murine K17 gain of function (GOF) cell line model. Murine Kras_G12D_, p53_R172H_ pancreatic cancer cells (KPC) barely express K17. Cells were transduced to stably express either EV (empty vector) or human K17 (K17) as K17 GOF models. Western blot and quantification are shown. **H.** Immunofluorescence images of KPC cell line model. Filament form of K17 is shown in green, and the nucleus is indicated by DAPI stain in blue. **I-J.** K17 causes chemoresistance to Gem (I) and 5-FU (J) in K17 GOF cell line models. K17-expressing KPC and MIA PaCa-2 cells had significantly higher IC50 values. IC50 values and fold change of IC50 are shown. **K-M.** Cell cycle analyses of K17 cell line models: L3.6 (K), MIA PaCa-2 (L) and KPC (M). **N-P.** Proliferation curves of K17 cell line models: L3.6 (N), MIA PaCa-2 (O) and KPC (P). No difference of proliferation between K17-positive and K17-negative cells was found. Data are shown in mean ± SD. *p<0.5, **p<0.01, ***p<0.001, ****p<0.0001, n=3-5. Student’s *t*-test.

We further validated these results in K17 gain-of-function (GOF) cell line models. Human PDAC cell line MIA PaCa-2 (Fig. 2E-F) and murine Kras_G12D_, p53_R172H_ pancreatic cancer cells (KPC Fig. 2G-H), which express low level of K17, were stably transduced to express K17 as K17 GOF cell line models. As proof of principle, K17-expressing cells showed significantly higher IC50 values compared to non-K17-expressing cells after treated with Gem (Fig. 2I, MIA PaCa-2: 10-fold, KPC: 5-fold) and 5-FU (Fig. 2J, MIA PaCa-2: 3-fold, KPC: 7-fold). Importantly, we did not detect any large difference in proliferation rates (Fig. 2N-P) or cell-cycle progression (Fig. 2K-M) between K17-positive and K17-negative cells in these three cell line models, suggesting that the differences in response to Gem and 5-FU are not related to differences in cell division. These findings indicate that K17 expression enhances the intrinsic resistance to chemotherapeutic agents in PDAC cells in a cell autonomous matter, suggesting that K17 could be used as a novel predictive biomarker in PDAC.

### 3.3. An unbiased high-throughput drug screen reveals compounds that specifically target K17-expressing PDAC

Considering that K17 expression drives chemoresistance to first-line therapeutic agents and that there are currently no therapeutic agents to target this intermediate filament, we set out to screen for small molecules that preferentially impact K17-expressing PDAC cells, with the goal of identifying therapeutic vulnerabilities in these cells and enhancing their response to current standard-of-care treatment.

We performed an unbiased high-throughput screen in isogenic L3.6 cells with or without expression of K17, against compounds from the Library of Pharmacologically Active Compounds (LOPAC) to identify the most effective and specific agents against K17-expressing cells (Fig. 3A). This primary screen yielded 24 active compounds (hits) that showed high response rates in L3.6 K17-expressing cells (Table 1). After completion of the first screen with K17-expressing cells, we performed a counterscreen assay with isogenic counterpart L3.6 K17 KO cells, and it yielded 16 active hits (Table 1). The hits from both the primary screen and counterscreen (p<0.0001) were then evaluated for overlap and grouped in three categories. 10 compounds were found to be selective for the K17-expressing cells, 14 compounds were found to equally target both K17-expressing and K17 KO cells, and 2 compounds were found to be selective for K17 KO cells (Fig. 3B, Table 1).

**Table 1.** Hits identified from the Screen and Counterscreen, with the cutoff parameter at 23% and 26%, respectively

| From | Drug Name | Description | Average Response Rate (%) |  |
| --- | --- | --- | --- | --- |
|  |  |  | K17 | K17 KO |
| S | Zardaverine | Phosphodiesterase III (PDE III) and phosphodiesterase IV (PDE IV) inhibitor | 46.69 | 26.00 |
| S | Vincristine sulfate | Inhibitor of microtubule assembly | 40.08 | 24.00 |
| S | Podophyllotoxin (PPT) | Inhibitor of microtubule assembly | 39.40 | 18.99 |
| S | Vinblastine sulfate salt | Inhibitor of microtubule assembly | 37.66 | 18.87 |
| S | Enoximone | Selective phosphodiesterase III (PDE III) inhibitor | 36.40 | 23.92 |
| S | DCEBIO | Increases epithelial chloride secretion through the synergistic activation of a basolateral membrane-located K<sup>+</sup> channel (hK1) and an apical membrane Cl<sup>-</sup> conductance | 32.95 | 16.96 |
| S | Diphenyleneiodonium chloride | Endothelial nitric oxide synthase inhibitor | 32.02 | 21.17 |
| S | Quazinone | Phosphodiesterase III (PDE III) inhibitor | 30.39 | 13.23 |
| S | PMA | Activates protein kinase C in vivo and in vitro; strong NO promoter; promotes expression of iNOS in cultured hepatocytes; T lymphocyte activator | 27.61 | 7.15 |
| S | Imazodan | Selective phosphodiesterase II (PDEII) inhibitor | 24.40 | 11.80 |
| S/C | Ouabain | Blocks movement of the H5 and H6 transmembrane domains of Na<sup>+</sup>-K<sup>+</sup> ATPases | 83.34 | 75.43 |
| S/C | Camptothecin | DNA topoisomerase I inhibitor | 69.58 | 70.77 |
| S/C | Mitoxantrone | DNA synthesis inhibitor | 65.63 | 71.47 |
| S/C | Emetine dihydrochloride hydrate | Apoptosis inducer; RNA-Protein translation inhibitor | 58.26 | 45.53 |
| S/C | Brefeldin A | Fungal metabolite that disrupts the structure and function of the Golgi apparatus | 57.72 | 45.18 |
| S/C | Dihydroouabain | Sodium-potassium pump inhibitor | 48.59 | 37.74 |
| S/C | Thapsigargin | Potent, cell-permeable, IP3-independent intracellular calcium releaser | 48.59 | 37.74 |
| S/C | Nocodazole | Disrupts microtubules by binding to beta-tubulin | 46.91 | 36.98 |
| S/C | Azathioprine | Purine analog; purine synthesis inhibitor; immunosuppressant | 44.85 | 52.99 |
| S/C | Cytarabine hydrochloride | Selective inhibitor of DNA synthesis | 33.90 | 36.36 |
| S/C | Ancitabine hydrochloride | Antineoplastic DNA metabolism inhibitor | 32.32 | 30.87 |
| S/C | Tyrphostin AG 1478 | Selective inhibitor of epidermal growth factor receptor (EGFR) protein | 30.67 | 32.03 |
| S/C | Paclitaxel (PTX) | Antitumor agent; promotes assembly of microtubules and inhibits tubulin disassembly process | 29.63 | 41.74 |
| S/C | 3-Amino-1-propanesulfonic acid sodium | GABA-A receptor agonist | 26.07 | 40.16 |
| C | N-(3,3-Diphenylpropyl)glycinamide | NMDA glutamate receptor open channel blocker. | 5.59 | 28.87 |
| C | Tyrphostin AG 879 | Tyrosine kinase nerve growth factor receptor (TrkA) inhibitor; inhibits 140 trk protooncogene and HER-2 | 6.28 | 28.94 |

S = Screen; C = Counterscreen

**Figure 3.**
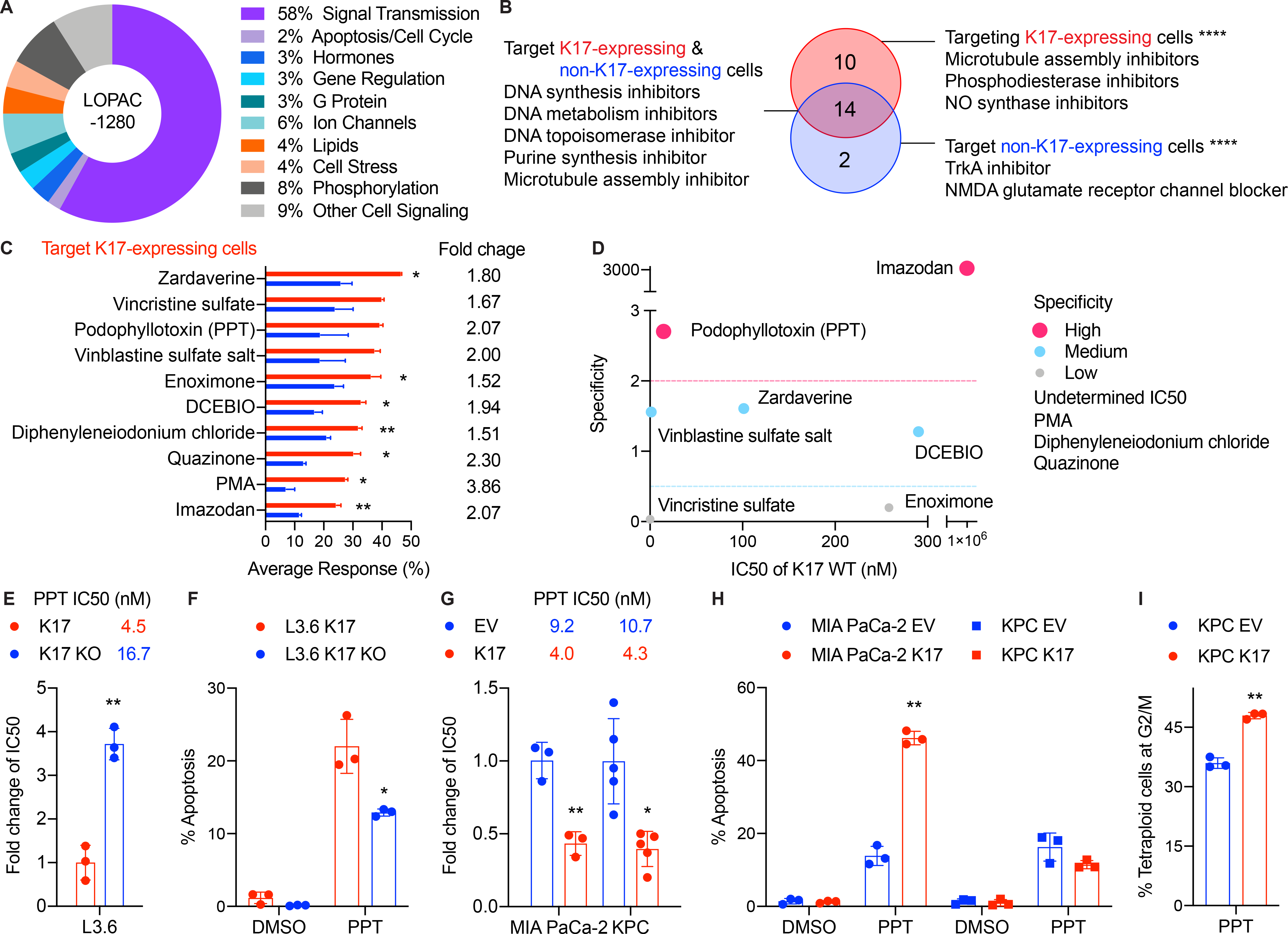
High-throughput drug screening identifies compounds that specifically target K17-expressing PDAC cells. **A.** LOPAC includes 1280 pharmacologically active compounds some of which are marketed drugs and pharmaceutically relevant structures annotated with various biological activities (shown in percent). **B.** The Screen (testing L3.6 K17-expressing cells) and the Counterscreen screen (testing L3.6 K17 KO cells) yielded 26 hits that were organized into three categories based on percent average response rate (at the p-values less than 0.0001). **C.** Drugs specifically targeting K17-expressing cells are listed. The average response rate and fold change of average response rate between K17-expressing and K17 KO cells are shown (mean ± SEM). **D.** Validation of the 10 drugs that be found to selective for K17-expressing cells. IC50 were determined in K17-expressing and K17 KO cells by CellTiter-Glo^®^ assay. The fold change of IC50 were calculated, and the specificity was then determined by the following equation: IC50 of K17 KO cells/IC50 of K17-expressing cells. Specificity, >2: high, 0.5-2: medium, <0.5: low. **F.** Validation of Podophyllotoxin (PPT) in L3.6 K17 LOF cell line model using WST-1 assay. K17 KO cells show significantly higher IC50 values than K17-expressing cells. IC50 values and fold change of IC50 are shown. **G.** L3.6 K17 KO cells showed significantly decreased percent apoptosis than K17-expressing cells under PPT treatment. **H.** Validation of PPT in K17 GOF cell line models using WST-1 assay. MIA PaCa-2 and KPC K17 cells showed significantly lower IC50 values than EV cells. IC50 values and fold change of IC50 are shown. **I.** MIA PaCa-2 K17 cells showed significantly increased percent apoptosis than EV cells under PPT treatment. No difference of percent apoptosis was found in KPC cells under PPT treatment. **J.** KPC K17 cells showed significantly higher cell population of tetraploid cells at G2/M phase than KPC EV cells under PPT treatment. Data are shown in mean ± SD. *p<0.5, **p<0.01, n=3-5. Student’s *t*-test.

The 10 identified compounds that show higher response rates in K17-expressing cells fell into two main categories: those that inhibit microtubule assembly (Vincristine sulfate, Podophyllotoxin [PPT] and Vinblastine sulfate salt) and those that inhibit phosphodiesterase activity (Zardaverine, Enoximone, Quazinone and Imazodan). In this LOPAC screen, three compounds, Zardaverine, Vincristine sulfate and PPT showed the highest response rates among the 10 candidate hits (Fig. 3C and Table 1). To validate the candidates, we determined the IC50 values in L3.6 cell line model and compounds were further defined as having high, medium or low specificity for K17-expressing cells (Fig. 3D). By cross-referencing results from these assays to determine selectivity, we found that PPT is the compound with the highest degree of specificity for K17-expressing cells and also required the lowest concentration to inhibit cell viability.

PPT is known to block cell division by destabilizing microtubule activity through binding tubulin^24^. To confirm the specificity of PPT in inhibiting K17-expressing PDAC cells, we performed an independent validation in additional K17-manipulated cell line models using the WST-1 cell viability assay. In L3.6 (Fig. 3E), MIA PaCa-2, and KPC (Fig. 3G) cell line models, PPT consistently demonstrated at least two-fold differences in the IC50 values of K17-expressing cells versus non-K17-expressing cells. Therefore, less than half of the dosage of PPT is required to exert the same inhibitory effect in K17-positive cells, as compared to K17-negative cells.

In addition, under PPT treatment, K17-positive cells showed significantly increased apoptosis compared to K17-negative cells in L3.6 (Fig. 3F) and MIA PaCa-2 (Fig. 3H) cells. In KPCs, a significantly increased population of tetraploid cells at G2/M phase were detected (Fig. 3I), as a result of the mechanism of action of this drug^24^; however, the proportion of apoptotic cells was similar between K17 and EV cells (Fig. 3H). Compared with additional validation experiments of Zardaverine from the same category (Fig. S1A and B), however, PTT was the only compound that consistently targeted K17-expressing cells across all models. Thus, through multiple independent validations of several cell lines, we identified that PPT is selectively effective in targeting K17-expressing PDAC cells.

Lastly, in the category of compounds that show similar efficacy in both 17-expressing and K17 KO cells, Mitoxantrone was selected for further studies because it shares a similar mechanism of action to current standard-of-care chemotherapeutic agents (DNA synthesis inhibitor) for PDAC^25^. Validation experiments showed that Mitoxantrone had similar sensitivity in K17-expressing and non-K17-expressing cells in all three cell line models (Fig. S1D-E). In addition, Tyrophostin AG 879, the drug targeting the K17 KO cells with the highest average response was chosen for further studies (Fig. S1H-J). We validated that this drug was much more sensitive in L3.6 K17 KO cells (Fig. S1I); however, it was highly resistant in MIA PaCa-2 and KPC cells (Fig. S1J), potentially due to phenotypic differences between cell line models or off-target effects. These verify results from the screen in L3.6 cell line model (Fig. S1C), and suggests that validation experiments of a high-throughput drug screen are necessary to confirm the on-target effect.

In summary, using an unbiased high-throughput drug screen, we identified and validated PPT as a potential drug to target K17-expressing PDAC cells.

### 3.4. Targeting microtubule disassembly rather than microtubule assembly is a therapeutic vulnerability in K17-expressing pancreatic cancers

To evaluate whether PPT might enhance Gem-mediated cytotoxicity and determine if the combined effect is synergistic, additive or antagonistic, we measured cell viability by treating with increasing doses of the combination of Gem and PPT at fixed ratios (1:1 in L3.6 cells and 37.5:1 in MIA PaCa-2 and KPC cells, based on their IC50 values of Gem and PPT, according to established protocols^26^). We found that K17-expressing cells had significantly lower cell viability under the same doses compared with non-K17-expressing cells (Fig. S2A-C). In addition, the IC50 of the combination of Gem and PPT was lower in K17-expressing cells compared to isogenic cells lacking K17 expression. We computed the combination index (CI)^23^ to determine the interaction of Gem and PPT in PDAC cells. Along with increasing effective doses from 50 to 97% of cell viability, the combination of Gem and PPT demonstrated a strongly synergistic interaction as shown by CI values lower than 0.9 in L3.6 K17-expressing cells, and there was an additive (effective dose at 50-75%) to antagonistic (effective dose at 90-97%) effect in L3.6 K17 KO cells (Fig. 4A). In MIA PaCa-2 cell line model, co-treatment showed an additive (effective dose at 50%) to antagonistic effect (effective doses at 75-97%) in K17-expressing (K17) cells and an antagonistic effect in non-K17-expressing (EV) cells (Fig. 4B). In KPC cells, a strongly synergistic effect was found in KPC K17 cells (CI<0.5) while an antagonistic effect was observed in KPC EV cells (Fig. 4C). In summary, we demonstrated that PPT in combination with Gem was synergistic or additive in K17-expressing cells but that these drugs had antagonistic effects in non-K17-expressing cells.

**Figure 4.**
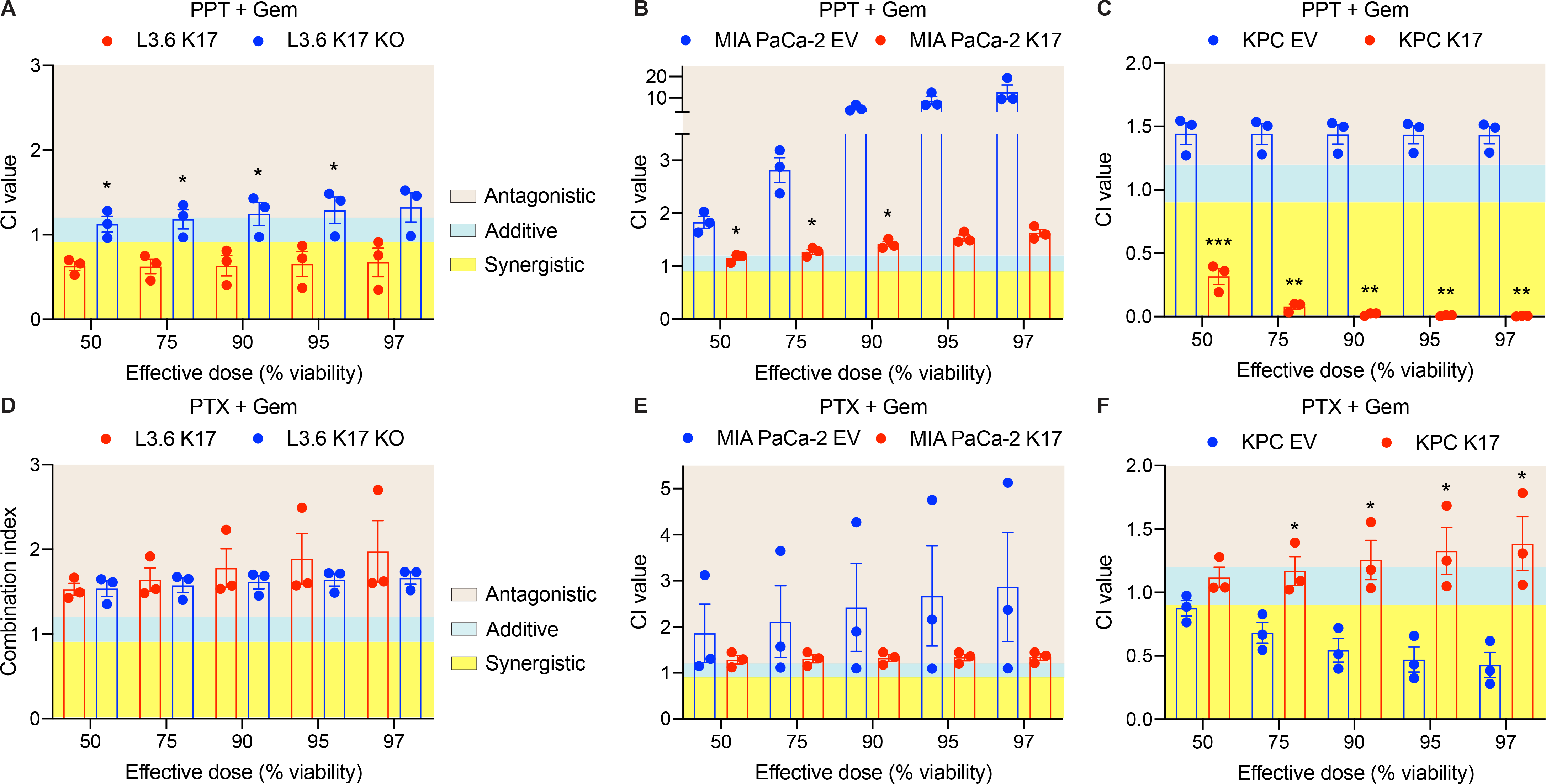
Targeting microtubule disassembly rather than microtubule assembly is a therapeutic vulnerability in K17-expressing pancreatic cancers. **A-C.** The combination effects of PPT and Gem in L3.6 (A), MIA PaCa-2 (B) and KPC (C) cell line models were determined by calculating the CI values at effective dose at 50, 75, 90, 95 and 97 percent of cell viability. **D-E.** The combination effects of PTX and Gem in L3.6 (D), MIA PaCa-2 (E) and KPC (F) cell line models were determined by calculating the CI values at effective dose at 50, 75, 90, 95 and 97 percent of cell viability. CI<0.9 indicates a synergistic effect. CI=0.9-1.2 indicates an additive effect. CI>1.2 indicates an antagonistic effect. Data are shown in mean ± SEM. *p<0.5, **p<0.01, ***p<0.001, n=3. Student’s *t*-test.

Currently, Gem is combined with Paclitaxel (Taxol, PTX), a compound that stabilizes microtubules and protects it from disassembly^27^ as a standard-of-care therapeutic regimen for PDAC^7^. Given that we identified PPT, a compound that destabilizes microtubules^24^, to be synergistic when combined with Gem in K17-expressing PDACs, we set out to determine whether the synergistic effect of Gem in combination with a microtubule inhibitor depends on a specific mechanism of action of PPT and/or PTX. First, we tested the IC50 values of PTX and found that they were similar in K17-positive versus K17-negative cells in L3.6 and KPC cell lines (Fig. S1F-G) and the IC50 was lower in K17 cells than in EV cells in MIA PaCa-2 cell line model (Fig. S1G). Second, we tested the combination of PTX and Gem, using the same experimental setup of testing PPT combined with Gem. We found no obvious differences in the dose-response curves of the combination in K17-positive and K17-negative cells (Fig. S2D-F). Surprisingly, we found that PTX and Gem tended to have antagonistic effects in K17-expressing cells across isogenic model systems (Fig. 4D-F), while it only showed synergistic effects in KPC cells lacking K17 expression (Fig. 4F).

Together, we found that K17 expression sensitizes cells to PPT in combination with Gem, however, these effects are adverse when Gem is combined with PTX in the same cells, suggesting that the mode of action of microtubule dynamic inhibitors may have a direct and/or indirect effect on K17 intermediate filament dynamic or function. Future studies, however, are needed to test this hypothesis.

## 4. DISCUSSION

Here, we identify that expression of K17 in PDACs correlates and causes resistance to Gem treatment, while it sensitizes cells to an FDA-approved compound that causes microtubule disassembly, identified by high-throughput screen and downstream validation assays. In addition, we found that the combination of Gem and this microtubule inhibitor synergizes to specifically target K17-expressing PDACs. This study is important as it reports a biomarker-based novel combination therapy that can be further tested in pre-clinical and clinical settings to target the most lethal molecular subtype of pancreatic cancer.

The report of the PDAC molecular subtypes was a key milestone for the advancement in understanding this highly lethal disease, opening three crucial research areas: 1) Development of clinical tests to subtype patients at the time of diagnosis; 2) Selection of the best standard of care (Gem/nab-PTX or FOLFIRINOX) for each molecular subtype; and 3) Identification of targeted therapies, based on PDAC subtypes.

It is critically important to develop clinical tests to subtype PDAC patients at the time of diagnosis, to provide the most appropriate therapy. We recently reported that K17 mRNA, a hallmark biomarker of the most lethal molecular subtype of PDAC, is as accurate as molecular subtyping to identify the PDAC patient population with the worst prognosis^15^. Moreover, K17 protein, as detected by immunohistochemistry (IHC), provides better prognostic value than mRNA. Importantly, we have established a protocol for K17 IHC that can be used both in research laboratory and in clinical laboratory diagnostic settings. As the only reported basal-like gene that provides both prognostic and predictive values, further studies are required to validate K17’s predictive value at the protein level using IHC.

To date, there are no predictive biomarkers to inform PDAC response to either Gem/nab-PTX or FOLFIRINOX regimens^28,29^, to enhance therapeutic efficacy, or to prevent adverse effects based on the molecular characteristics of the tumors. Based on our findings, K17 may be a candidate predictive biomarker for response to Gem/nab-PTX. Our data also suggests that K17 promotes resistance to 5-FU, a main agent in FOLFIRINOX. As such, it is important to determine if K17 is a predictive marker of FOLFIRINOX in PDAC. Importantly, the COMPASS trial findings show that basal-like PDACs tended to be more resistant to the FOLFIRINOX chemotherapy^17^. To our knowledge, K17 is the only gene expressed in the most lethal basal-like subtype of PDAC that actively promotes chemoresistance to Gem and 5-FU and thus, may represent a potential clinical target for personalized treatment.

With the goal of identifying novel targeted therapies using existing FDA-approved compounds that can be repurposed and fast-tracked for the treatment of the patients with K17-expressing PDACs, we performed a high-throughput drug screen using the LOPAC assay. Although previous LOPAC screens had been applied to PDAC cell lines^30^, to date, these screens have not been evaluated in the context PDAC molecular subtypes and in combination with current chemotherapeutic agents. Through validation studies in three different K17 loss- and gain-of-function PDAC cell line models, we report that Podophyllotoxin (PPT) was consistently found to preferentially target K17-expressing cells. Our findings suggest that microtubule assembly inhibitors may be potentially effective drugs for treating K17-expressing PDACs. These observations are novel because the combination of PPT and Gem as a potential regimen for cancer therapy has not been previously reported. Furthermore, we also found that PPT was the most effective agent to target K17-expressing PDAC cells and that it synergizes in combination with Gem. Surprisingly, these effects were not recapitulated when combining Gem with PTX. This suggests that the current standard-of-care therapy may be replaced by a more effective and synergistic combination: Gem and PPT. Although beyond the scope of this study, future studies should address the mechanistic basis regarding how K17 expression sensitizes PDAC cells to PPT treatment.

Importantly, compared to other microtubule inhibitors from our LOPAC screen (such as PTX, Vincristine sulfate and Vinblastine sulfate salt), PPT showed the highest specificity in inhibiting K17-expressing PDACs. In a previous study, PPT was shown to significantly inhibit the growth of lung tumor cells and using *in vitro* and *in vivo* models, was cytotoxic in several human cancer cell lines^31–34^. Although PPT is currently only used for treatment of HPV-mediated cutaneous lesions, its semisynthetic compounds, Etoposide and Teniposide, are widely used for cancer therapy in the clinic. A phase II trial previously showed that Gem combined with Etoposide exhibited a response rate similar to published trials using Gem-based chemotherapies^35^. Despite extensive interest in utilizing PPT and its derivates for cancer therapy, very few of these agents have reached clinical practice, in part due to toxicity and solubility^36–38^. More selective PPT derivatives based on modified structures may enhance effective responses in K17-expressing PDACs. Future studies of testing the specificity of Etoposide and Tenitoposide, alone and in combination with Gem in K17-expressing PDACs are needed.

In summary, we discovered a novel therapeutic vulnerability of K17-expressing PDACs and identified a compound that when combined with Gem, may enhance tumor response, compared to the current standard-of-care with Gem/nab-PTX.

## 5. Conclusions

The identification of biomarker-driven novel therapies is a critical priority for improving survival of PDAC patients. K17 is a novel target for development of a biomarker-based personalized treatment for the most aggressive form of PDAC. We demonstrated that beyond its predictive value, K17 drives chemoresistance to Gem and 5-FU. Through an unbiased drug screen, we discovered that PPT showed at least 2-fold higher sensitivity in K17-positive PDAC cells. Surprisingly, we found that small molecules that inhibits microtubule disassembly (PPT) rather than microtubule assembly (PTX) specifically targets K17 PDACs, uncovering a novel therapeutic vulnerability of the most aggressive molecular subtype of PDAC. These studies serve as scientific premise to further launch pre-clinical testing of this novel combination therapy that can guide optimal therapeutic management, which is driven by data related to the molecular composition and biomarker expression in PDACs.

## Data accessibility

Not applicable.

## Author contributions

C-HP, TS, LFE-H and KRS designed the research. SB, DKC and AB performed patient data analyses. C-HP, CVL and JDB generated cell line models. YO and BS performed drug screening. C-HP, YO and MW performed validation experiments and analyzed data. LS and TS supervised the high-throughput compound screening study. LFE-H and KRS supervised the patient data analyses and *in vitro* validation study. C-HP and MW wrote the manuscript. YO, CVL, TS, LFE-H and KRS edited and revised the manuscript. All authors have read and approved the final manuscript.

## Acknowledgements

The authors thank Dr. Peter Bailey and the APGI for RNAseq data, Dr. Coulombe Pierre (University of Michigan) for providing the K17 antibody, Dr. Wei-Xing Zong (Rutgers University) for L3.6 cells, Dr. Gerardo Mackenzie (University of California at San Diego) for KPC cells and all members of our laboratory for critical review of the manuscript.

---

**Figure S1.**
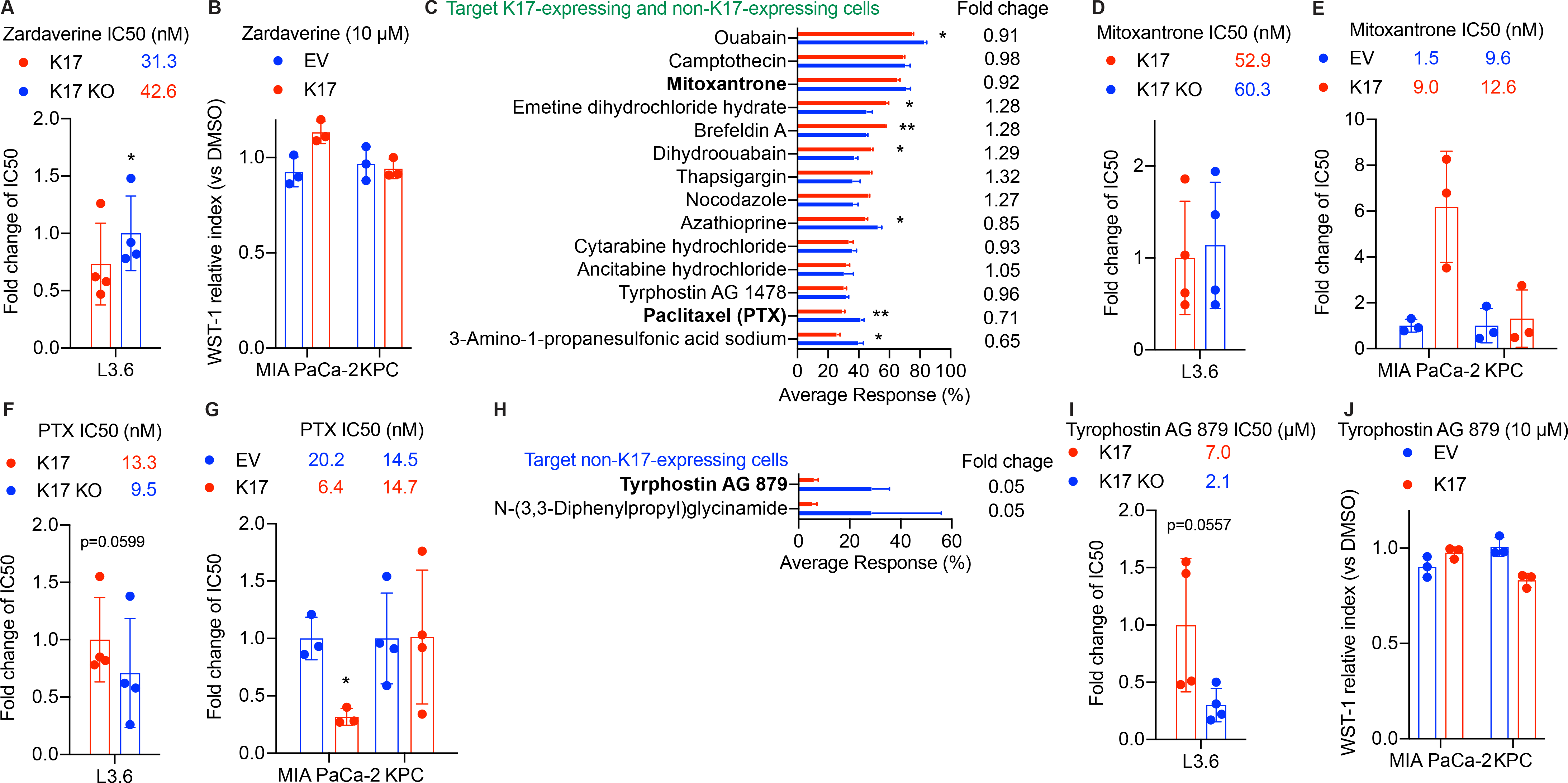
Validation of compounds in other categories. **A-B.** Validation of Zardaverine in L3.6 (A) and in MIA PaCa-2 and KPC cell line models (B). IC50 values, fold change of IC50 or cell viability (WST-1 relative index) are shown. **C.** Drugs targeting both L3.6 K17-expressing and KO cells from the Screen and the Counterscreen are listed. Fold change of average response rate are shown (mean ± SEM). **D-E.** Validation of Mitoxantrone in L3.6 (D) and in MIA PaCa-2 and KPC cell line models (E). IC50 values and fold change of IC50 are shown. **F-G.** Validation of PTX in L3.6 (F) and in MIA PaCa-2 and KPC cell line models (G). IC50 values and fold change of IC50 are shown. **H.** Drugs targeting L3.6 K17 KO cells from the Screen and the Counterscreen are listed. Fold change of average response rate are shown (mean ± SEM). **I-J.** Validation of Tyrophostin AG879 in L3.6 (I) and in MIA PaCa-2 and KPC cell line models (J). IC50 values, fold change of IC50 or cell viability (WST-1 relative index) are shown. Data are shown in mean ± SD. *p<0.5, **p<0.01, n=3-4. Student’s *t*-test.

**Figure S2.**
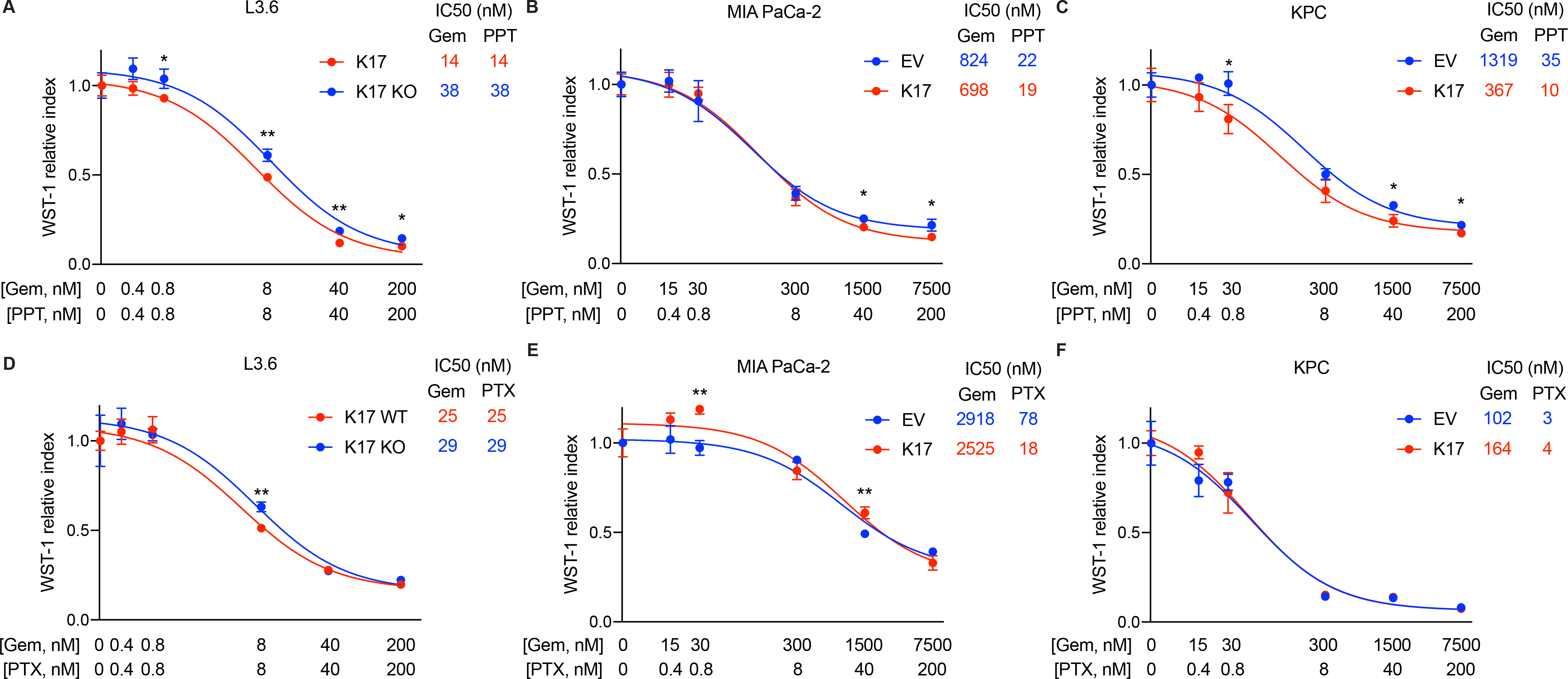
K17-expressing cells show lower cell viability than non-K17-expressing cells under treatment of PPT and Gem, but no obvious difference is found in PTX and Gem. **A-C.** The dose-response curves of PPT combined with Gem were shown in L3.6 (A), MIA PaCa-2 (B) and KPC (C) cell line models. The predicted IC50 of PPT + gem in each cell line were listed. **D-F.** The dose-response curves of PTX combined with Gem were shown in L3.6 (D), MIA PaCa-2 (E) and KPC (F) cell line models. The predicted IC50 of PPT + gem in each cell line were listed. Data are shown in mean ± SD. *p<0.5, **p<0.01, ***p<0.001, n=3. Student’s *t*-test.

